# Multisensory perception reflects individual differences in processing temporal correlations

**DOI:** 10.1101/264457

**Authors:** Aaron R. Nidiffer, Adele Diederich, Ramnarayan Ramachandran, Mark T. Wallace

**Affiliations:** Department of Hearing and Speech Sciences, Vanderbilt University, Nashville, TN, USA; Department of Health, Life Sciences & Chemistry Jacobs University, Bremen, Germany; Vanderbilt Brain Institute, Vanderbilt University, Nashville, TN, USA; Department of Psychology, Vanderbilt University, Nashville, TN, USA; Department of Psychiatry, Vanderbilt University, Nashville, TN, USA; Vanderbilt Kennedy Center, Vanderbilt University, Nashville, TN, USA

**Author notes:** Corresponding Author: Aaron Nidiffer 7110 MRB III BioSci Bldg 465 21^st^ Avenue South Nashville, TN 37240-7933.

**Keywords:** audiovisual, binding, decision making, diffusion model

## Abstract

Sensory signals originating from a single event, such as audiovisual speech, are temporally correlated. Correlated signals are known to facilitate multisensory integration and binding. We sought to further elucidate the nature of this relationship, hypothesizing that multisensory perception will vary with the strength of audiovisual correlation. Human participants detected near-threshold amplitude modulations in auditory and/or visual stimuli. During audiovisual trials, the frequency and phase of auditory modulations were varied, producing signals with a range of correlations. After accounting for individual differences which likely reflect relative temporal processing abilities of participants’ auditory and visual systems, we found that multisensory perception varied linearly with strength of correlation. Diffusion modelling confirmed this and revealed that correlation is supplied to the decisional system as sensory evidence. These data implicate correlation as an important cue in audiovisual feature integration and binding and suggest correlational strength as an important factor for flexibility in these processes.

## Introduction

Our environment provides us with an enormous amount of information that is encoded by multiple sensory modalities. One of the fundamental tasks of the brain is to construct an accurate and unified representation of our environment from this rich array of sensory signals. To accomplish this, the brain must decide which signals arise from a common source. For example, during conversation among a group of individuals, listeners can group appropriate words from the same voice and further associate voices with the appropriate speakers, a process greatly facilitated by the availability of both audible and visible cues (Stein 2012; M. Murray and Wallace 2012). Benefits that are associated with the presence of multisensory signals include increased detection (Frassinetti, Bolognini, and Làdavas 2002; Lovelace, Stein, and Wallace 2003) and localization accuracy (Ohshiro, Angelaki, and DeAngelis 2011; Stein, Scott Huneycutt, and Alex Meredith 1988), speech intelligibility (Sumby and Pollack 1954; Erber 1969) and speeding of reaction times (Hershenson 1962; Hughes et al. 1994; Harrington and Peck 1998; Frens, Van Opstal, and Van der Willigen 1995; Corneil et al. 2002).

In realistic and complex sensory environments, neural signals within and across the different sensory systems that relate to an object must be integrated appropriately while also being segregated from signals associated with unrelated events. The “binding problem” (A. Treisman 1998, 1996) concerns how the brain achieves a coherent representation of the environment by deciding elements of the world belong together. Several mechanisms have been proposed that attempt to explain this process (Barlow 1972; A. M. Treisman et al. 1980; Olshausen, Anderson, and Van Essen 1993; Singer and Gray 1995), including synchronized activity across brain regions resulting from similarity in the temporal features of an object (Singer and Gray 1995; Engel, Senkowski, and Schneider 2007; Senkowski et al. 2008). Multisensory signals originating from the same source, such as the voice and mouth movements of a speaker, are temporally correlated (Chandrasekaran et al. 2009). Temporal correlation has been shown to be a robust cue for the binding of unisensory (Fahle 1993; Blake 2005; Elhilali et al. 2009; Micheyl, Hanson, et al. 2013; Yost and Sheft 1994) and multisensory (Vatakis and Spence 2007; Parise, Spence, and Ernst 2012; Parise et al. 2013; Munhall et al. 1996) features. Human observers can utilize temporal correlations in multisensory signals, resulting in enhancements in behavioral performance compared to uncorrelated multisensory signals (Maddox et al. 2015; Grant and Seitz 2000). Further, a number of human multisensory behaviors can be adequately described using a “correlation detection” model focused on the correlation between signals from the different sensory modalities (Parise and Ernst 2016).

Although we know that temporal correlation between unisensory signals leads to a unified multisensory percept and enhancement of multisensory behaviors, it is not known whether, and if so how, multisensory behavioral performance varies proportional to the strength of the correlation. We hypothesize that audiovisual temporal correlation provides sensory evidence as to whether stimulus features belong to the same event, and that this evidence is proportional to the direction and strength of the correlation. Further we hypothesized that these graded changes in sensory evidence result in corresponding changes in multisensory behavior. To test this, we presented participants with audiovisual signals with barely detectable amplitude modulation (AM). While manipulating the temporal correlation between the auditory and visual AM, we measured how observers’ ability to detect these fluctuations changed with changes in stimulus correlation. We propose a mechanism—analogous to a phase shift—that approximates relative differences in unisensory temporal processing and that accounts for individual differences in behavioral results. Finally, we employed drift-diffusion modelling to test whether multisensory behavioral performance is better approximated by absolute stimulus correlation or by adjusted correlations that account for this phase shift.

**Figure 1.**
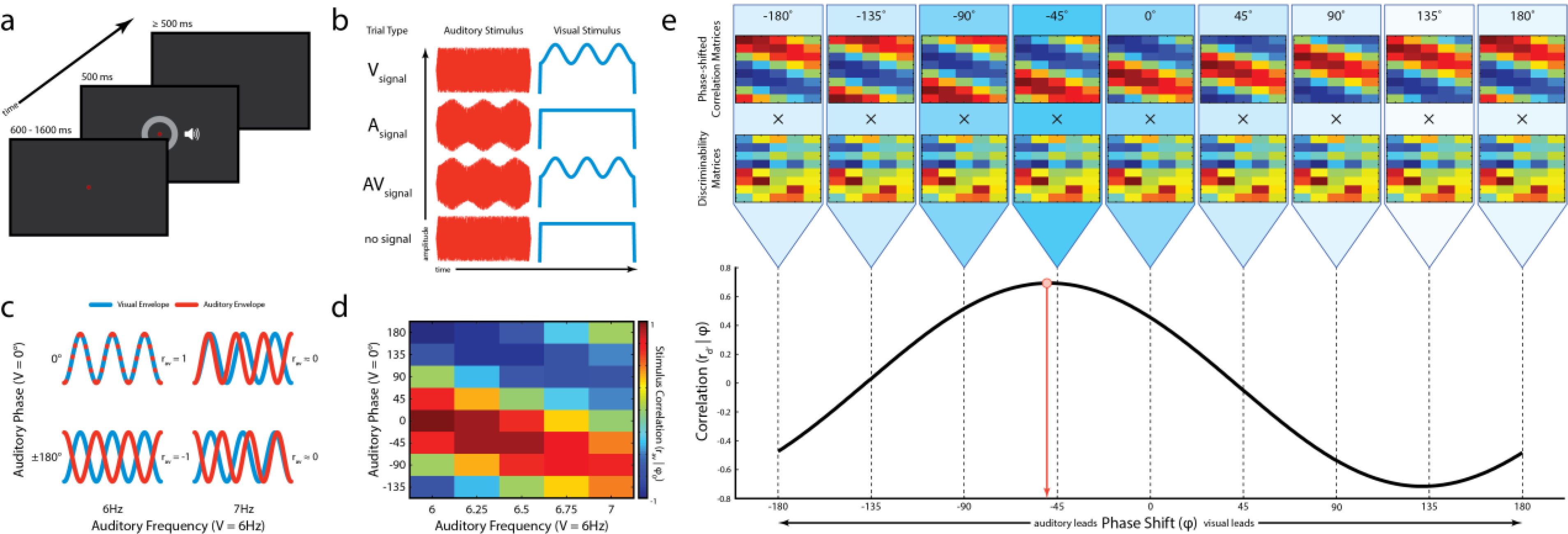
Amplitude modulation detection task. **a**: Schematic representation of a single trial. Each trial began with the illumination of a fixation target. After a variable wait period, simultaneously presented auditory and visual stimuli appeared (see b). Participants indicated the presence or absence of amplitude modulation with a button press. **b**: Auditory and visual stimuli were always present, but modulation was presented in auditory stimuli only (Asignal trial), visual stimuli only (Vsignal trial), audiovisual stimuli (AVsignal trial), or neither stimulus (no signal, catch trial). **c**: During audiovisual presentations, the frequency and phase of auditory modulation could be independently manipulated yielding a range of audiovisual correlations (r_av_). Correlations were computed using the time series of the auditory and visual envelopes. Note that the visual envelope is always constant while the auditory envelope is varied. Four conditions out of forty are shown for illustration. **d**: Stimulus Correlation Matrix (r_av_ | ϕ_0_). All forty AV stimulus conditions are shown organized according to Δ frequency × Δ phase. Colors represent the correlation values of audiovisual stimuli across the different frequencies and phases presented where each color box represents one condition. In the task structure, there were 21 unique audiovisual stimulus correlations. **e**: In order to account for phase shifts in individual participant data, the values in the stimulus correlation matrix (r_av_ | ϕ_0_) were correlated to each participant’s discriminability matrix (r_d’_). In the top panel, a series of correlation matrices are shown in which a phase lag, ϕi, was applied to auditory (positive shifts) or visual (negative shifts) before correlations were computed, (r_av_ | ϕi). Each correlation matrix was correlated with the participant’s discriminability matrix (r_d’_; middle panel), approximating a cross correlation. In the bottom panel, these correlations (r_d’_) were plotted against phase lag [(r_d’_ | ϕ); black line]. This function was fit to a sine wave and the phase of that fit was extracted (ϕ’; red dot and arrow) and was taken to represent a participant’s individual phase shift. The stimulus correlation at that phase shift (r_av_ | ϕ’) was taken to represent a participant’s “internal” correlation matrix.

## Results

Participants detected near-threshold amplitude modulated (AM) audiovisual stimuli where the temporal correlation of the AM signals was manipulated by systematic changes in the phase and frequency relationship of the auditory and visual pairs. Our central hypothesis was that multisensory behavioral performance would improve commensurate with increasing temporal similarity between the paired audiovisual stimuli (i.e., as correlation become more positive). To examine the potential dependence of behavior on stimulus correlation, a discriminability (*d’*) matrix and a reaction time (RT) matrix for each participant was constructed and related to the stimulus correlation (r_av_) matrix (Δ frequency × Δ phase; Figure 1d).

While RTs did not show any systematic pattern, discriminability had a discernible pattern relative to the stimulus correlation matrix. In eight of 12 participants, discriminability was significantly correlated with stimulus correlation (Figure 2a; sig. r = 0.64 ± 0.15 [mean ± standard deviation]). However, upon visual inspection, the discriminability matrices of two of the remaining four participants mirrored the stimulus correlation matrix but with an apparent shift along the Δ phase dimension (see Figure 2a-b, middle panels for one example). In fact, this phase shift appeared to be present in most participants to varying degrees and seemed to occur evenly across Δ frequency for each participant (i.e., any shift along the phase dimension was present for all auditory frequencies presented). We therefore hypothesized that this phase shift reflects an internal transformation that alters the relationship between stimulus correlation and behavior.

**Figure 2.**
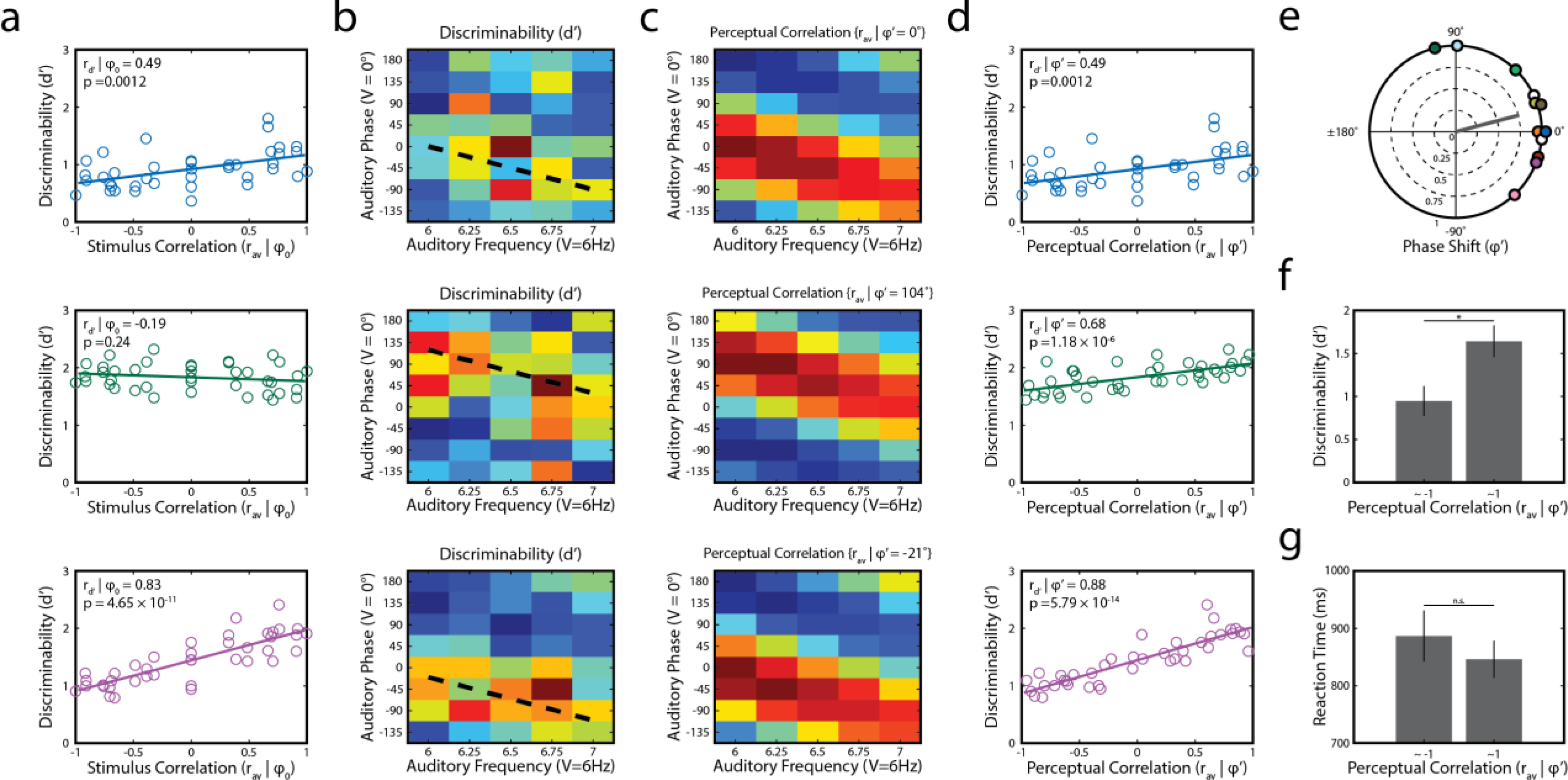
Individual participant data examples. **a**: Behavioral dependence on stimulus correlation (r_d’_ | ϕ_0_) of three example participants. For parts a-d, each row represents a single participant. Colors follow the convention described in Figure 3a. **b**: Discriminability matrices from three participants plotting how changes in phase (@del phase; y-axis) and frequency (@del frequency; x-axis) impact the ability to detect amplitude modulation (discriminability). Diagonal dashed lines represent the computed individual phase shifts (ϕ’ = 0°, −104°, and 21°) corresponding to the approximate middle of the diagonal of positive correlations in (c). **c**: Phase shifted (“perceived”) correlation matrices (r_av_ | ϕ’) from each participant shown in (a). Note the strong positive (upward) shift in the second example participant and the moderate negative (downward) shift in the third example participant, relative to figure 1d. **d**: Behavioral dependence (r_d’_ | ϕ’) on perceived stimulus correlation. Each participant shows a strong positive relationship between perceived stimulus correlation and detection behavior (i.e., discriminability). Note that the data in the middle panel was not significantly correlated to physical stimulus correlation (a) but reached significance when accounting for the phase shift. Colors follow the convention described in Figure 3a. **e**: Distribution of observed phase shifts from all participants and mean resultant vector. Phase shifts were concentrated around the mean (14.7°, not uniform across phase). Phases were shifted toward positive values (visual leading), but were not significantly different from zero. **f-g**: Accuracy and reaction time effects between stimuli with the strongest negative and positive perceptual correlations. Strong positive correlation improves detection performance, but has no impact on reaction times.

### Individuals display unique characteristics for auditory and visual temporal processing

We sought to measure and account for these individualized phase shifts. We hypothesized that unisensory temporal processing differences were the reason behind these shifts. We modeled this by applying a phase shift to every condition in one of the unisensory modalities before recalculating a stimulus correlation matrix. We then measured the correlation between the discriminability matrix and a series of stimulus correlation matrices computed with phase shifts ranging from −180° to +180° (Figure 1e; more detail in methods). We then fit this series of correlations to a sine wave. Due to the cyclical nature of the stimulus correlation matrix along the Δ phase dimension, we expected the correlations to be in the shape of a sine wave. As expected, each participant’s phase-shifted correlations were well fit (r^2^ = 0.99999 ± 2.9 × 10^−5^). Another expectation is for these functions to have a period of 360° and to be centered about zero, which we found to be the case (period: 360.06 ± 0.71; *t_11_* = 0.2702, *p* = 0.79; center: 1.3 × 10^−4^ ± 5.4 × 10^−4^; *t_11_* = 0.783, p = 0.45). Therefore, we calculated a participant’s phase shift from these fits and then recomputed a unique correlation matrix for each participant using their individual phase shift.

As a test of the validity of phase shift, the pattern of data in the discriminability matrix should mirror the pattern of the phase-shifted stimulus matrix. This would manifest in several ways. First, if the perceived correlation matrix accounts for the data, large changes in the data should be accounted for by changes in the correlations. Therefore, the residual errors between the two measures should be very small relative to the data and centered around zero. Discriminability values (Figure 2b) were significantly above zero (*d’* = 1.30 ± 0.66; *z* = 43.579, *p* = 8.75 ×10^−169^). Subtracting the predicted *d’*, which were computed from the perceptual correlation matrices (see Methods; Figure 2c), from the observed *d’*, yielded residual errors which were substantially smaller and less variable compared to *d’* (mean error = 0.018 ± 0.33). Indeed, these residual errors did not differ significantly from zero (*z* = 1.210, *p* = 0.23). Second, we might question the validity of these phase shifts if the data do not mirror perceptual correlations equally for each Δ frequency (e.g., if the diagonal of high *d’* values in the discriminability matrix has a slope that doesn’t match the slope of strong correlation values in the correlation matrix). To quantify this, we examined residual errors across different frequencies for any systematic changes. Residual error magnitude and variability showed no linear relationships across Δ frequency in any participant (magnitude: slopes = 0.047 ± 0.10, all *p* > 0.12; variability: slopes = 0.016 ± 0.07, all *p* > 0.09). Thus, phase shifts appear to be valid and systematic shifts in the phase dimension and are independent of frequency. As such, the correlation matrices constructed using each participant’s unique phase shift could be envisioned to represent the internal (“perceived”) correlations of the external stimuli, accounting for differences in latency of sensory processing between the auditory and visual systems.

These perceptual correlations were used when determining the relationship between discriminability and stimulus correlation (r_d’_; Figure 2d). The sine wave fits between phase shift and correlation revealed the degree of participant audiovisual phase shift (ϕ’; Figure 2e). Phase shifts were not significantly different from 0 across participants but favored a visual leading shift (mean ϕ’ = 14.7 ± 39.7°; 95% CI [42.2° −12.9°]). The distribution of shifts was concentrated about the mean as indexed by the mean resultant vector length (Figure 2e; MRVL = 0.76; z = 11.998, p = 1.5×10^−8^, Rayleigh Test). To further probe the validity of these phase shifts, we tested whether the magnitude of phase shift was correlated to the strength of the relationship between behavior and stimulus correlation. Smaller correlations associated with larger phase shifts might suggest that the repeated phase shift approach returned spurious correlations. We found no evidence of such a relationship (rho = 0.25, p = 0.68). *Amplitude modulation discriminability varies with perceived stimulus correlation*

Previously, it has been shown that strongly correlated multisensory stimuli provided behavioral and perceptual benefits relative to unisensory performance whereas poorly correlated stimuli fail to provide such benefits (Maddox et al. 2015; Parise et al. 2013; Parise, Spence, and Ernst 2012). To examine whether a similar relationship is evident for the current task, we compared the discriminability of stimuli that had the highest and lowest correlation for each participant. We found that discriminability of audiovisual signals with the highest correlations was better than for audiovisual signals with the lowest correlations (Figure 2f; t11 = 4.312, p = 0.0062, corrected). In contrast, reaction times failed to differ between correlated signals and uncorrelated signals (Figure 2g; t11 = 3.384, p = 0.19, corrected).

Our focus of the current study was to show that multisensory behavior varied proportionally with stimulus correlation. Although we demonstrated above that this relationship was robust in most participants (Figure 2a), there was evidence that this effect was weakened—and in some participants absent—due to significant individual variability. Thus, it still remained unclear whether phase shift plays an important role in this relationship. To test this, we measured the association between perceived stimulus correlation and discriminability (r_d’_ | ϕ’; Figure 3a). These correlations were significant in ten out of the twelve participants—two participants more than when not accounting for phase shift. This proportion, 10/12, was significantly greater than expected based on random effects, 0.05 (*p* = 0.019, binomial test). The significant correlations revealed effects that were very strong (sig. r = 0.72 ± 0.14, all sig. r > 0.49, all p < 0.0012; Figure 3b). Indeed, when accounting for phase shift, the strength of these correlations increased in all participants (Δr_d’_ = 0.19 ± 0.29; sig. Δr_d’_ = 0.22 ± 0.31) and the increase was more pronounced with larger magnitude phase shifts (Figure 3e, aobs = 0.706). Due to the nature of the phase-shift fitting process, simulated random data (details can be found in methods) produces correlational improvement that peaks at ± 180° (a = 0.205, 95% CI [0.144 0.271]). Nonetheless, the observed effect was significantly larger than would be expected by these random effects (z = 21.80, p = 2.1×10^−93^). Lastly, in contrast to the concentrated distribution of observed phase shifts (Figure 2e), the distribution of simulated phase shifts was not significantly different from uniform (Figure 3f; MRVL = 0.04; z = 2.08, p = 0.125, Rayleigh Test). These findings provide strong support for the notion that phase shift reflects an important transformation between stimulus correlation as it occurs in the environment and how it manifests in perceptual performance.

**Figure 3.**
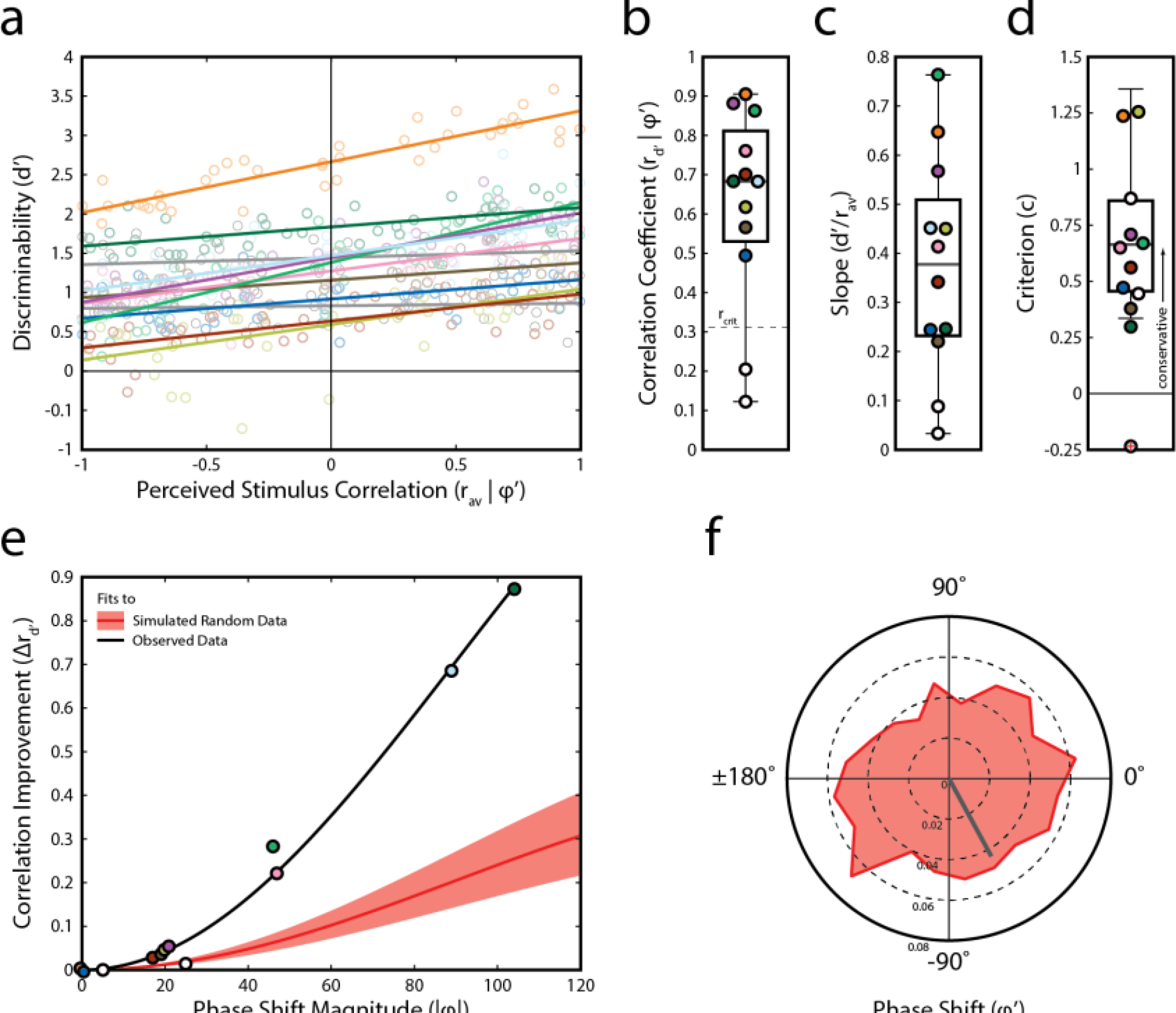
Behavioral results. **a**: Behavioral dependence on perceived stimulus correlation across all participants (same as Figure 2d). Behavioral performance in 10 of 12 participants was driven by stimulus correlation. Non-significantly correlated data are represented in grey. Significantly correlated data is depicted in color, with each participant retaining the same color throughout the figures. **b**: Correlation coefficients (r_d’_ | ϕ’) for each participant. Colors follow the convention described in (a). **c**: Slope of linear data fits shown in (a) for each participant. Colors follow the convention described in (a). **d**: Criterion for each participant. Each participant but one held a conservative criterion indicating that participants weren’t biased toward responding “yes.” Colors follow the convention described in (a). **e**: Improvement in correlation (Δr_d’_) is associated with phase shift and the effect is larger than expected by chance. Red line and shaded region represent the average fit and 99% confidence bands of random data from the Monte Carlo simulation fit to a sine wave. The black line represents the fit of the observed data to the sine wave. The amplitude of the data fit sine wave was significantly larger than expected by chance. Colors follow the convention described in (a). **f**: Distribution of phase shifts and the corresponding MRVL obtained from the Monte Carlo simulation. In contrast to observed data shown in Figure 2e, these phase shifts are not significantly concentrated about the circular mean. Note the scale difference in the radial axis between Figure 2e and here.

Individuals showed widely varying dependencies on stimulus correlation as measured by the slope of a linear psychometric function fit to discriminability data (Figure 3c; sig. slopes = 0.43 ± 0.18). Lastly, despite the stimuli being presented at threshold levels, we were concerned about the possibility of participants adopting a strategy that exploits our low proportion of catch trials (i.e., they could be always reporting the presence of the stimulus modulation). We therefore quantified the participant willingness to respond with “modulation present.” Figure 3d confirms that this strategy was not employed by participants (c = 0.61 ± 0.41) with 11 of 12 adopting a conservative criterion. Further reinforcing this, 10 out of 12 participants (including the lone participant with a liberal bias) were within one standard deviation of an unbiased criterion (−1 < c < 1).

### Perceived stimulus correlation predicts audiovisual behavior via changes in evidence accumulation

Next, we sought to describe how audiovisual temporal correlation and phase shift influence behavioral performance in a decisional framework. Typically, changes in choice frequency and reaction time in a decision task are driven by changes in sensory evidence. We hypothesized that, in our task, sensory evidence was conferred by the temporal correlation of the stimuli. Further, we asked whether perceptual correlations rather than physical correlations may better account for changes in behavioral performance in a subject-by-subject basis. To answer these questions, we employed two decision models that were constrained to test these hypotheses. The first model assumed that the drift rates, which index sensory evidence, are related to physical stimulus correlations (r_av_ | ϕ0) across conditions. For model 2 we assumed that the drift rates are related to the perceived stimulus correlations, that is, correlations determined after a phase shift was applied (r_av_ | *ϕ_i_*). This design allowed the models not only to predict choice and reaction times with sensory evidence based on stimulus correlation, but also to measure participant phase shifts, providing converging evidence (in conjunction with results provided above) of an internal phase shift of the representation of the physical stimuli.

**Table 1.** Model 1 parameters.

| Ptc. | $\theta$ | $\beta$ | $T_r$ | $w$ | $X^2$ | AIC |
| --- | --- | --- | --- | --- | --- | --- |
| 1 | 13 | 0.2151 | 0.5741 | 0.0182 | 97.719 | -0.281 |
| 2 | 7 | 0.4954 | 0.7915 | 0.0762 | 87.192 | -10.808 |
| 3 | 19 | 2.3893 | 0.6113 | 0.0111 | 175.113 | 77.113 |
| 4 | 9 | 3.5568 | 0.7605 | 0.0001 | 70.945 | -27.055 |
| **5** | **13** | **-4.4554** | **0.8** | **0.0018** | **125.043** | **27.043** |
| 6 | 9 | 6.2835 | 0.6364 | 0.0001 | 95.408 | -2.592 |
| 7 | 8 | -0.6953 | 0.6018 | 0.0334 | 142.682 | 44.682 |
| 8 | 15 | 0.1974 | 0.788 | 0.0292 | 108.722 | 10.722 |
| 9 | 17 | 0.1539 | 0.7956 | 0.025 | 131.553 | 33.553 |
| 10 | 5 | 1.3622 | 0.7738 | 0.0339 | 88.424 | -9.576 |
| 11 | 16 | -11.9243 | 0.588 | 0.0333 | 163.444 | 65.444 |
| **12** | **9** | **3.2324** | **0.7994** | **0.0089** | **142.744** | **44.744** |

**Table 2.** Model 2 parameters.

| Ptc. | $\Phi$ | $\theta$ | $\beta$ | $T_r$ | $w$ | $X^2$ | AIC |
| --- | --- | --- | --- | --- | --- | --- | --- |
| 1 | 46 | 16 | 1.3753 | 0.4859 | 0.0214 | 214.55 | 14.55 |
| 2 | 0 | 7 | -0.2467 | 0.7881 | 0.078 | 158.4 | -41.6 |
| 3 | 0 | 19 | 2.3988 | 0.6074 | 0.011 | 281.68 | 81.681 |
| 4 | -104 | 6 | 2.6569 | 0.7915 | 0.0348 | 136.03 | -63.97 |
| **5** | **-2** | **13** | **-4.4437** | **0.8** | **0.0012** | **217.53** | **17.528** |
| 6 | -92 | 8 | 4.6167 | 0.6443 | 0.0519 | 181.62 | -18.382 |
| 7 | 13 | 10 | -2.0867 | 0.5639 | 0.0298 | 252.04 | 52.041 |
| 8 | 19 | 15 | 0 | 0.7863 | 0.0317 | 168.74 | -31.256 |
| 9 | -46 | 17 | -1.3041 | 0.8 | 0.0374 | 208.79 | 8.79 |
| 10 | -16 | 14 | 2.3778 | 0.6013 | 0.0133 | 237.15 | 37.146 |
| 11 | -8 | 15 | -11.4521 | 0.6079 | 0.0375 | 225.2 | 25.201 |
| **12** | **2** | **9** | **2.2282** | **0.7951** | **0.0086** | **242.31** | **42.309** |

Table 1 and 2 show the estimated parameters for each model and their goodness of fit. Both models were well fit to the data and model 2 successfully incorporated the extra parameter for phase shift without compensation from other parameters meant to index bias, decision style (conservative vs. liberal), and sensory encoding/preparation. As evidence that the models were not simply adjusting other parameters to adjust between models, we found that these parameters were strongly correlated between models when accounting for phase shift using partial correlations (θ: rho = 0.78, p = 0.0046; β: rho = 0.98, p = 7.67 × 10^−8^; Tr: rho = 0.87, p = 0.00044). Using Akaike Information Criterion (AIC) as a model selection metric, we found that most participants’ behavior was better described by the second model, in which the perceived correlation, included as a phase shift parameter, drives the decision process. Qualitatively, perceptual choice across conditions can be described as a dampening oscillator with dampening increasing with Δ frequency, a pattern which is also apparent in the model prediction of choice. Figure 4a shows the model fit (colored lines matching conditions shown in Figure 4b) to a single participant’s data (filled circles).

**Figure 4.**
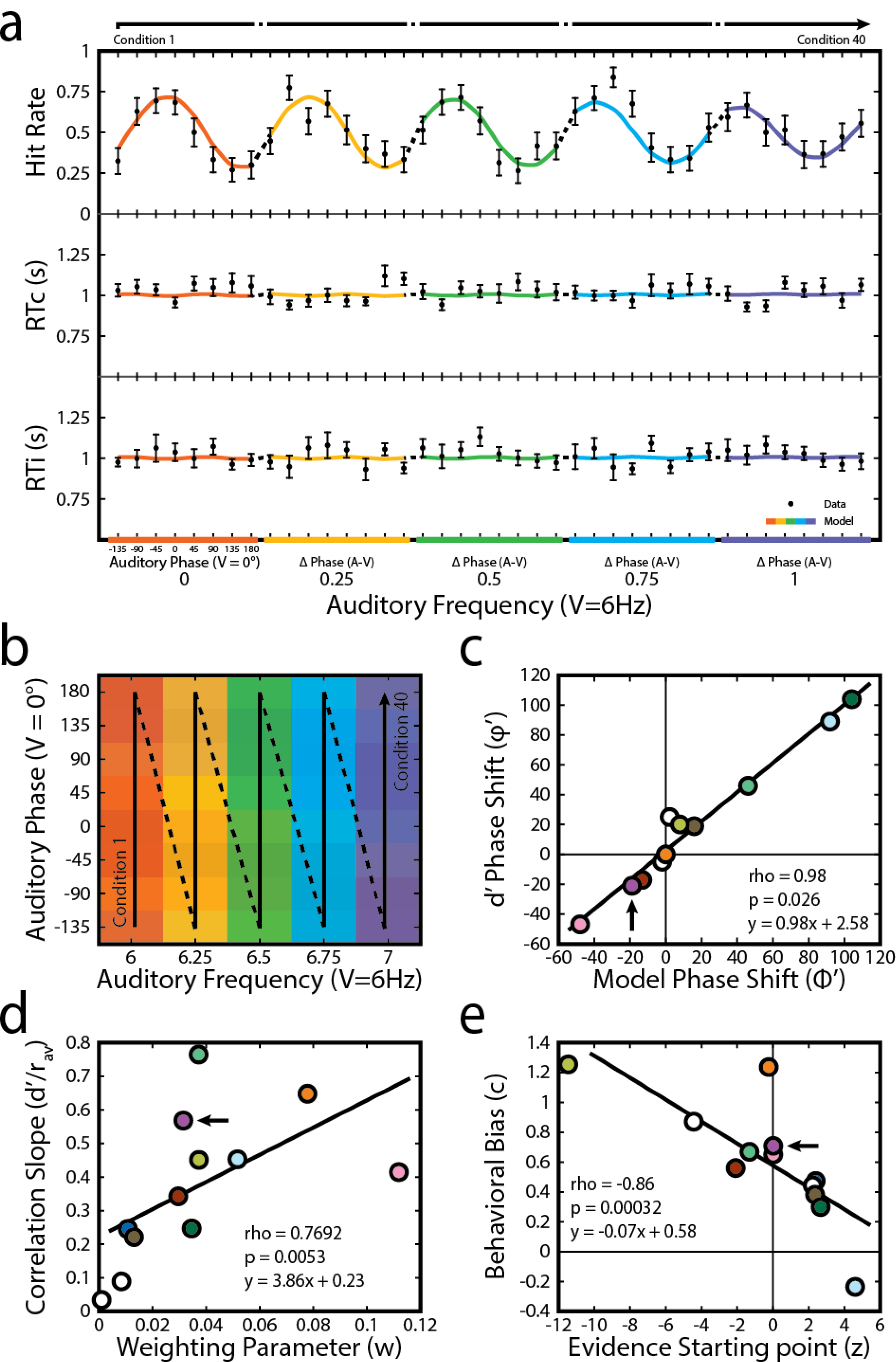
Modeling results and comparison to behavioral results. **a**: Model fit for a single participant. Proportion of correct responses (top panel), reaction times for correct responses (RTc; middle panel), and reaction times for incorrect responses (RTi; bottom panel) are shown (black dots, ± 1 S.E.M) for all 40 audiovisual conditions (top arrow). The same data are shown from the model prediction (colored lines). Model predictions and observed data are shown along a single continuous axis for simplicity with non-continuous data points connected by dashed lines (see panel b for key). **b**: Stimulus correlation and discriminability are organized in matrices (Figure 2b-c) with columns representing different Δ Frequencies and rows representing different Δ Phases. In (a), data have been reorganized column-wise such that Condition 1 is the first Δ Phase in the first Δ Frequency and Condition 40 is the last Δ Phase in the last Δ Frequency. Colors of the model fit and bottom axis in (a) correspond to columns in the matrix with the same color. Solid portions of the top arrow in (a) correspond to vertical solid portions of the arrow in (b). Similarly, dashed portions of the arrow in (a) correspond to oblique dashed portions in (b). **c**: Across all participants, phase shifts measured from discriminability matrices (Figure 2e) are strongly correlated with the phase shift parameters output by the diffusion model. Participant data shown in (a) correspond to the marker indicated by the arrow. Colors follow the convention described in Figure 3a. **d**: Measures of bias, criteria (from Figure 3d) and evidence starting point parameters are correlated across participants. Participant data shown in (a) correspond to the marker indicated by the arrow. Colors follow the convention described in Figure 3a. **e**: Measures of dependence on stimulus correlation, psychometric slopes (from Figure 3c) and scaling parameters are correlated across participants. Participant data shown in (a) correspond to the marker indicated by the arrow. Colors follow the convention described in Figure 3a.

Model 2 made accurate predictions of behavioral choice and reaction times based on the perceptual correlations returning parameters that closely matched their signal detection theory counterpart. Each participant’s model-fit phase shift parameter (*ϕ’*) nearly perfectly matched their phase shift obtained from discriminability (*ϕ’*, Figure 4c; rho = 0.98, p = 0.026, slope = 0.98). Additionally, evidence starting point, *β*, which is the parameter that measures the participant’s bias toward one response over another (Voss, Rothermund, and Voss 2004; Laming 1968), was also correlated with the signal detection theory measure of bias, *c* (Figure 4d; rho = 0.77, p = 0.0053). The bias reflects the participant’s tendency to respond with “modulation present” or “modulation absent”, which is unrelated to the sensitivity of the participant. Lastly, the drift-rate weighting coefficient was strongly linked correlated with the slope of their psychometric functions (Figure 4e; rho = −0.86, p = 0.00032), with both measures describing the dependence of behavior on changes in correlation. Moreover, the parameters were very consistent between models.

## Discussion

Temporal factors such as (a)synchrony have long been known to influence multisensory processes in the brain (Meredith, Nemitz, and Stein 1987; Wallace, Wilkinson, and Stein 1996; Bushara, Grafman, and Hallett 2001; E. Macaluso, Frith, and Driver 2002; Emiliano Macaluso et al. 2004; Senkowski et al. 2007) and in behavior (Hershenson 1962; Frens, Van Opstal, and Van der Willigen 1995; Dixon and Spitz 1980; McGrath and Summerfield 1985; Stone et al. 2001; Colonius and Diederich 2004; Fujisaki et al. 2004). More recently, Parise and colleagues (2012) presented evidence that the fine temporal structure of an audiovisual stimulus *independent of asynchrony* can influence multisensory perception. They further showed that it is possible to explain a number of multisensory phenomena based on a general correlation detection mechanism (Parise and Ernst 2016).The findings presented in the current study provide unique but additional support for the growing evidence implicating temporal correlation as an important cue in multisensory processing.

In the current work we extend this knowledge about multisensory temporal dependencies by showing that audiovisual detection behavior is a monotonic function of stimulus correlation. As the temporal similarity of two unisensory signals increased, detection of amplitude modulation embedded in the audiovisual signal improved in a linear manner (Figure 3a). Additionally, we qualify this finding in a way that provides mechanistic insight into how the brain combines dynamic stimuli across sensory modalities. Thus, the temporal correlation of the audiovisual stimuli did not necessarily map directly onto multisensory behavioral performance; conditions in which physical stimulus correlation was highest did not always result in the best behavioral performance. Instead, it appears that a transform occurs in the brain of each individual and that results in a phase shift in behavioral performance relative to physical stimulus correlation (r_av_ | *ϕ_0_*). Calculating temporal correlation after applying a phase lag to one of the stimuli (r_av_ | *ϕ’*), which simulates different processing times of the sensory signals in the brain, accounts for this difference. These phase-shifted correlations presumably represent the correlations as they are available to our decisional system.

Although our task did not reveal any measurable effects of temporal correlation on reaction times, we are not surprised. This lack of effect can be explained in terms of RT variability. Our stimuli employed near-threshold signals which are known to produce reaction times that are more variable than produced by supra-threshold signals (McKendrick, Denniss, and Turpin 2014). Thus, our reaction times might have too much variability to measure any meaningful effect. Additionally, the correlations in our stimuli unfold over time. For some conditions (*f_m, auditory_* = 6 Hz), the correlation doesn’t change throughout the course of the signals. However, for others they change over time. This difference could introduce more reaction-time variability in some conditions than in others and thus mask a potential effect. To measure any potential effect in reaction times, an experiment would need to be designed using supra-threshold signals. *Temporal correlation as a unit of sensory evidence*

The current study places the relationship between stimulus correlation and multisensory processing in a decisional framework. Our model successfully incorporated the relationship between two signals (i.e., temporal correlation) into a dynamic-stochastic approach to account for choice frequency and response time. With only very few parameters (4 for model 1 and 5 for model 2) stimulus correlation was able to account for the observed patterns. Moreover, it was able to account for individual differences within and across participants. Our primary finding is related to the nature of how stimulus correlation influences the accumulation of sensory evidence for a decision. Specifically, we found that perceived (phase-shifted) stimulus correlation serves as a good predictor of behavior when used to constrain drift rate. For perceptual tasks, drift rate is often interpreted as an index for the quality (e.g., strength) of sensory evidence that is available to the decisional system (Gold and Shadlen 2007; Ratcliff and Smith 2004). Typically, the strength of sensory evidence is provided by the physical attributes of the stimulus, for instance, the degree of motion coherence (Ratcliff and McKoon 2008), intensity (Ratcliff 2002; Rach, Diederich, and Colonius 2011), line length (Adele Diederich and Busemeyer 2006; Adele Diederich 2008), or numerosity (Leite 2012) in discrimination tasks. For simple multisensory behaviors (e.g., detection of simple stimuli), the drift rate relates to the combined evidence obtained from integrating the physical stimulus properties across modalities (Rach, Diederich, and Colonius 2011; Schwartz 1989; A Diederich and Colonius 1991; Schwartz 1994; a Diederich 1995; Otto and Mamassian 2012), especially when these properties are weak or ambiguous (e.g., low intensity, poor motion coherence, etc.) in the unisensory component stimuli (Rach, Diederich, and Colonius 2011).

In the current task, the key physical parameter that would presumably modulate the magnitude of evidence for detection is the depth of the amplitude modulation, with strength of evidence increasing with depth. However, modulation depth, and thus sensory evidence from the unisensory signals, is held constant across conditions. Although we cannot rule out that evidence is supplied by integration of the unisensory stimulus properties, sensory evidence cannot come from these alone but instead is generated during a computation involving both stimuli. Different types of multisensory decisions require different architectures that depend on the structure of the task or stimulus (Bizley, Jones, and Town 2016). The results presented here—that the strength of sensory evidence is based on a computation of the unisensory signals rather than the strength of the unisensory signals themselves—suggests that unisensory signals converge and evidence is computed prior to being evaluated by the decisional system. Other multisensory decisions such as simultaneity judgement (Simon, Nidiffer, and Wallace, n.d.) and temporal order judgement (Mégevand et al. 2013; Adele Diederich and Colonius 2015), which require a comparison of the unisensory signals, have also been described in terms of a cross-modal computation.

### Does binding occur proportional to the strength of temporal correlation?

It has been well established that temporal correlation plays a role in binding unisensory features in both auditory (Elhilali et al. 2009; Micheyl, Hanson, et al. 2013; Yost and Sheft 1994) and visual (Fahle 1993; Blake 2005) domains. For example, in an auditory grouping task, two tones, A and B, are repeated sequentially: ABABAB… If the difference in their frequencies becomes sufficiently large, the two tones are segregated and perceived as two separate streams rather than a single “trill” stream. However, if the temporal relationship is manipulated such that A and B occur at the same time, perceptual fusion is re-established (Micheyl, Kreft, et al. 2013). More recently, it has been shown that temporal correlation between auditory and visual signals results in their binding (Parise et al. 2013; Parise, Spence, and Ernst 2012; Maddox et al. 2015). The result of the binding of correlated audiovisual signals manifests in a variety of ways, many of which have been demonstrated with speech signals. The intelligibility of speech in a noisy environment is improved with the addition of visual information (Sumby and Pollack 1954). This improvement requires that modulations of lip movements to be temporally correlated with the modulations of the speech envelope (Munhall et al. 1996; Grant and Seitz 2000), as is the case in natural audiovisual speech (Chandrasekaran et al. 2009). This effect is thought to be mediated by an enhanced cortical representation of auditory speech that is dependent on correlation between auditory and visual speech streams (Crosse, Butler, and Lalor 2015). Results presented here support this concept, despite the substantially different nature of the stimuli employed.

According to our results, multisensory benefits—and likely the propensity to bind two signals—are monotonically related to the strength and sign of the temporal correlation (similarity) between unisensory signals. This notion implies that the process of binding signals is probabilistic at some level. Stochastic binding related to temporal correlation could be an important mechanism in cognitive flexibility. In a sensory-rich environment, compulsory binding based on temporal similarity could lead to unification of unrelated stimuli. Instead, binding and integration relies on several features such as spatial and temporal coincidence. In the natural environment, these features are very often in agreement; a single event will produce energies across different modalities that overlap in space and time and that are temporally correlated. Where these features are somewhat discrepant, the brain will appropriately weight the features in the construction of a multisensory percept (Ernst and Banks 2002). For example, when the locations of auditory and visual signals are ambiguous both in absolute and relative terms, but their onsets are synchronous, the brain often combines them despite their spatial discrepancy (Alais and Burr 2004). Similarly, the brain loses information about the relative spatial location of temporally correlated audiovisual stimuli (Parise et al. 2013). The linear relationship between stimulus correlation and perception could represent a weighting cue for whether two signals should be bound. However, the linear nature of these effects may not hold for all stimulus or tasks (Parise et al. 2013).

### Entrained oscillations could explain the link between stimulus correlation and behavior

The perceptual benefits of increased stimulus correlation could be attributed to mechanisms involving synchronized or coherent neural activity across brain regions. Neural coherence has been hypothesized to play a role in shaping our conscious experience (Tononi and Koch 2008) by underpinning mechanisms of sensory awareness (Engel and Singer 2001; Melloni et al. 2007), attentional selection (Schroeder and Lakatos 2008), cognitive flexibility (Fries 2005; Womelsdorf et al. 2007), and perceptual binding (Senkowski et al. 2008; Singer and Gray 1995; Elhilali et al. 2009). Rhythmic stimuli like the ones used in the current study are known to entrain neural oscillations (Thut, Schyns, and Gross 2011) which index patterns of neuronal excitability over time (Bishop 1933). Temporally congruent audiovisual stimuli cause an increase in the coherence of neural activity *within* their respective unisensory sensory cortices (Nozaradan, Peretz, and Mouraux 2012). Finally, coherence *across* brain networks has been shown to vary with perceptual experience associated with multisensory binding (Hipp, Engel, and Siegel 2011).

In addition to promoting binding of audiovisual stimuli, temporal correlation has also been implicated in improved unisensory processing. Temporally congruent auditory and visual streams result in more inter-trial phase coherence in auditory and visual cortices (Nozaradan, Peretz, and Mouraux 2012). In auditory cortex of ferret, neuronal encoding of an auditory stream is improved by the presentation of a temporally congruent visual stimulus (with the same amplitude envelope) but not an incongruent stimulus (Atilgan et al. 2017). Further, the presence of a congruent visual stream improved the neuronal encoding of an orthogonal auditory feature (e.g., timbre shifts) embedded in the stimulus, which has been argued to be a hallmark of audiovisual binding (Bizley, Maddox, and Lee 2016). In human listeners this results in improved perception of those orthogonal auditory features (Maddox et al. 2015). Additionally, discrimination of the spatial features (orientation) of a temporally dynamic visual stimulus can be improved by the presentation of a temporally congruent auditory stimulus (Kösem and van Wassenhove 2012). Coupled with the finding that rhythmic unisensory features in multiple frequency bands entrain oscillations that interact to influence perception of orthogonal features (Molly J Henry, Herrmann, and Obleser 2014), it follows that the degree of temporal correlation of the amplitude envelope of audiovisual stimuli as were used in the current study would influence perception of other (e.g., spectral) features of the stimuli in a similar manner. This is likely the mechanism by which visible facial movements improves speech intelligibility (Sumby and Pollack 1954; Erber 1969; Grant and Seitz 2000).

### Differences in A and V entrainment could account for behavioral phase shifts

In the current study, participants’ behavioral performance was not necessarily best for the stimuli with highest physical correlation but were instead phase-shifted by differing amounts for each participant. Behavior very closely matched the correlation of the modulations after a phase lag was applied to one of the modulation signals. This phase lag could be adjusting for different processing times and abilities of participants’ auditory and visual systems. It’s known that oscillations entrain to rhythmic auditory stimuli at different phase lags across listeners (M. J. Henry and Obleser 2012). It is possible that visual entrainment occurs in a similar manner and that these phase lags differ between the auditory and visual systems. Interestingly, phase lag of the entrained oscillations can be calibrated to the particular temporal structure of an audiovisual stimulus (Kösem, Gramfort, and Van Wassenhove 2014). Thus, the phase lags reported in the current study are likely a “preferred” or “natural” phase that can be easily manipulated depending on context (e.g., attending an event that is near or far from the body which would result in different temporal relationships between auditory and visual representations in the brain) in a manner similar to the phenomenon of recalibration of the perception of audiovisual simultaneity (Fujisaki et al. 2004; Van der Burg, Alais, and Cass 2013).

### Conclusion

During multisensory decisions, temporal correlation between the features of the component stimuli modulates behavior. It does so by changing the nature of the sensory evidence that is evaluated by the sensory system. The strength of the sensory evidence is proportional to the strength of the correlation of the signal. Finally, the physical correlations present in stimuli are transformed, via a phase shift, into “perceptual” correlations that are unique to an individual. This process likely occurs through differences in unisensory temporal processing. This was confirmed by a dynamic-stochastic model in which the drift rate was related to physical or to perceived correlations between the auditory and visual signals in the audiovisual presentation. These results motivate several new questions. 1. Is binding truly stochastic? Can cross-modal correlation embedded in one feature (e.g., intensity) have the same proportional effect on behavioral performance reported here in tasks utilizing orthogonal stimulus features (e.g., timbre)? 2. What are the neural underpinnings of this proportional change and their relation to differences in unisensory processing? 3. Does the perception of natural stimuli such as speech benefit proportionally with correlation?

## Materials and Methods

### Participants

Twelve individuals (age = 26.4 ± 5.1, seven females) participated in the current study. All participants reported normal or corrected-to-normal vision and normal hearing and were right handed. The study was conducted in accordance with the declaration of Helsinki, and informed written consent was obtained from all participants. All procedures were approved by the Vanderbilt University Institutional Review Board. When applicable, participants were given monetary compensation for participation.

### Apparatus and stimuli

All stimuli were generated in MATLAB (The MathWorks, Inc., Natick, MA) and presented using PsychToolbox version 3 (Brainard 1997; Kleiner et al. 2007). Auditory stimuli were digitized at 44.1 kHz, and presented through calibrated open-back circumaural headphones (Sennheisser HD480). Visual stimuli were centered about a red fixation dot in the center of a dark (0.15 cd/m^2^) viewing screen (Samsung Sync Master 2233rz, 120 Hz refresh rate).

Auditory stimuli were frozen tokens of white noise (generated by the *randn* function) at moderate level (48 dB SPL, A-weighted). Visual stimuli consisted of a moderately bright ring (24 cd/m^2^; inner diameter: 1.8°, outer diameter: 3.6° visual angle). Both stimuli were presented simultaneously, lasted 500 ms, and were gated by a linear 10 ms onset and offset ramp. Stimulus timing was confirmed with a Hameg 507 oscilloscope, photodiode, and microphone.

Each stimulus, *y*, was modulated over time, *t*, such that

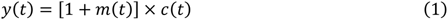

where

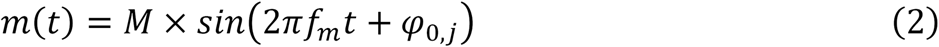

and *c(t)* is the time series of the carrier stimulus (auditory: noise; visual: ring). The form of the amplitude modulation (AM) signal *m(t)* is defined by a modulation depth *M* which represents the amplitude of the modulation signal as a proportion of the amplitude of the carrier signal and ranged from 0 (no AM) to 1 (full AM), frequency *f_m_* in Hz, and starting phase *ϕ_0,j_* in degrees.

On any given trial, the AM signal could be present in the auditory channel alone, the visual channel alone, both channels (audiovisual trials), or neither (catch trials; Figure 1b). Unisensory signals (AM was present in auditory stimulus only or visual stimulus only) were always presented in cosine phase such that the modulation began at the trough (*ϕ* = 0°) and at the same frequency (*f_m_, visual* = 6 Hz). When AM was present in both stimuli, visual modulation was always 6 Hz and cosine starting phase while auditory signals could be presented at various frequencies (*f_m_, auditory* = {6, 6.25, 6.5, 6.75, 7 Hz}) and initial phases (*ϕ_0_* = {−135, −90, −45, 0, 45, 90, 135, 180°}, with *ϕ_0,j_* ∊ ϕ_0_). This structure results in a total of 40 (5 × 8) different audiovisual stimulus conditions.

Because we are interested in the temporal correlation between the two signals, the Pearson correlation between the auditory and visual envelopes (r_av_) was computed for each of the 40 audiovisual conditions (Figure 1c). For example, when the auditory and visual envelopes were characterized by the same frequency and phase, correlation was 1. Conversely, stimuli of the same frequency but presented anti-phase resulted in a correlation of −1. The parameters chosen resulted in a representation of correlations between −1 and 1. A stimulus correlation matrix (r_av_ | ϕ_0_) was constructed for all audiovisual conditions by organizing the correlation values according to their frequency and phase relationship between auditory and visual signals (Δ frequency × Δ phase; Figure 1d).

### Procedure

Participants were seated comfortably inside an unlit WhisperRoom™ (SE 2000 Series) with their forehead placed against a HeadSpot™ (University of Houston Optometry) with the forehead rest locked in place such that a participant’s primary eye position was centered with respect to the fixation point at the center of the viewing screen. Chinrest height and chair height were adjusted to the comfort of the participant.

Prior to the main experiment, each participant completed two separate 3-down 1-up staircase procedures to obtain 79.4% modulation depth thresholds for auditory and visual AM at 6 Hz. For these staircase procedures, on a given trial (Figure 1a), the red fixation dot appeared at the center of the screen. Participants were instructed to fixate the dot for its entire duration. After a variable time, either an auditory or visual stimulus was presented in which the presence of modulation was determined at random for each trial. Participants were instructed to report the presence of amplitude modulation (described as “flutter”) after the stimulus presentation by pressing “1” on the number pad of a computer keyboard if the modulation was present or pressing “0” if the modulation was absent. The modulation depth decreased after three successive correct responses and increased after one incorrect response. At the beginning of each staircase, the step size was set to increase or decrease modulation depth by 0.05. After two reversals (correct to incorrect response or incorrect to correct response), step size was reduced to 0.025. Finally, after eight reversals, step size became 0.01 in order to arrive at an accurate estimate of modulation depth threshold. Each staircase terminated after 20 reversals. Threshold was determined to be the average of the modulation depth at the last 10 reversals. Instructions included an example of a stimulus with AM at the initial starting modulation depth (*M = 0.5*) and an example of a stimulus with no AM. So that there was no ambiguity in cases where the first trial did not include a modulation signal, participants were informed that the first trial would have the same modulation depth as the example if present. To control for “runs” of trials with no modulation during the staircase (which could result in erroneously low threshold estimates), a sequence of two trials containing no modulation was always followed by a trial with modulation. The auditory staircase was always completed first and served as a period of dark adaptation prior to the visual staircase.

The main experiment consisted of four blocks lasting approximately 30 minutes each. Each block consisted of 10 trials of each stimulus condition (420 signal trials per block). Additionally, there were catch (no signal) trials included to make up 10% of total trials for that block (47 catch trials per block). Therefore, each block was identical in trial composition (467 total trials per block) but with individual trials presented in a predetermined, pseudorandom order. Each participant completed a total of 1868 trials over the four blocks. Breaks were offered frequently (every 100 trials) to prevent fatigue. Participants completed the full experiment in 2-4 sessions, never completing more than 2 blocks during a session. If a participant completed two blocks in a single session, they were given the opportunity to stretch and walk around while the experimenter set up the second block. Before each block and after any break where the participant was exposed to normal light levels, participants were dark adapted for five minutes. Trials during the main experiment were identical to staircase trials with three exceptions. First, in each trial, both auditory and visual stimuli were presented. Modulation signals could be present in the visual channel alone (V_signal_), auditory channel alone (A_signal_), in both (AV_signal_; with frequency and phase configuration discussed above), or neither channel (no signal). Second, modulation depth was set to a participant’s unique auditory and visual modulation depth thresholds. Last, participants were told that they should respond as soon as they had made their decision and were instructed to respond as quickly and accurately as possible. In addition to the participant’s choice, response times were recorded for each trial, sampling every 2.2 μs (4.6 kHz).

### Behavioral Analysis

Discriminability (*d’*; a measure of sensitivity) for each of the 40 audiovisual conditions and two unisensory conditions was computed from the relative frequencies of the respective responses,

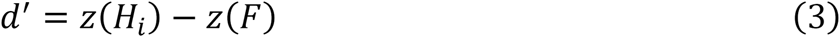

where *H_i_* is the proportion of hits (”1” | modulated stimulus) for the i^th^ condition, *F* is the proportion of false alarms (”1” | no modulated stimulus), and *z* is the inverse of the normal distribution function (MATLAB’s *norminv* function) and converts the hit rates and false alarm rates into units of standard deviation of a standard normal distribution. *d’* was organized into a matrix in the same manner as the stimulus correlation matrix. Because the proportion of catch trials was held low and errors had no associated cost (Green and Swets 1966), participants could potentially adopt a strategy of simply pressing “1” which would result in a correct choice more often than not. To account for this, criterion (*c*; a measure of bias) for each participant was computed in a similar manner such that

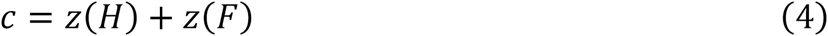

where *H* is the proportion of hits across all conditions. A single criterion was computed for each participant.

To account for individual differences, which became apparent in assessing the phase shift in the *d’* matrices, a series of correlation matrices based on the stimulus correlation matrix (r_av_ | ϕ_0_) were computed after iteratively applying a single degree phase lag to one stimulus (i.e., *ϕ_1_* = {−134, −89, −44, 1, 46, 91, 136, −179°}, *ϕ_2_* = {−133, −88, −43, 2, 47, 92, 137, −178°}, in general *ϕ_i_* = {−135 + *i*, −90 + *i*, −45 + *i*, 0 + *i*, 45 + *i*, 90 + *i*, 135 + *i*, 180 + *i*} with *i* = −180, …, 180, resulting in a total of 360 different matrices). A phase-shifted correlation matrix (r_av_ | *ϕ_i_*) could be conceptualized as the “internal” or “perceived” correlation of the signals given a particular phase lag, *i*, of one of the signals. Each of the phase-shifted correlation matrices (Figure 1e, nine examples shown) was in turn evaluated for correlation (r_d’_) with the discriminability matrix of each participant. The resulting correlation values (r_d’_ | ϕ) were then fit to a sine wave using the nonlinear least-squares method. The phase shift value of the fitted sine wave was recorded for each participant (ϕ’). The CircStat toolbox(Berens 2009) was used to describe the nature of the phase shifts and compute the directional statistics across the sample of participants. The “perceptual” correlation matrix corresponding to each participant’s unique phase shift (r_av_ | ϕ’) was used to measure the dependence of behavior on perceived correlation (r_d’_ | ϕ’).

To show that phase shift is related to a central mechanism (e.g., a relative difference in processing latencies between auditory and visual systems), we tested whether the phase shift occurred systematically across all Δ frequencies within each participant. First, a predicted discriminability matrix was calculated from phase-shifted correlations. Phase-shifted correlation matrices were normalized to each participant’s discriminability range by scaling and shifting each unique correlation matrix such that the correlation values at the maximum and minimum correlation matched the *d’* values at the corresponding locations in the discriminability matrix. Next, the values in the predicted discriminability matrix were subtracted from the actual discriminability matrix, resulting in a matrix of residual errors. Then, a linear model was used to determine the relationship (i.e., slope) between Δ frequency and the magnitude and variability (standard deviation) of errors. To calculate significance of variability slope across Δ frequency, a permutation test was used that shuffled the Δ frequency label of errors before calculating standard deviation within each Δ frequency and then fitting a line to the shuffled standard deviations.

We sought to demonstrate that accounting for phase shift improved the measured correlation between behavior and stimulus correlation. Therefore, we computed this dependence on stimulus correlation (r_d’_ | ϕ_0_) and subtracted it from the dependence on perceived correlation discussed above (r_d’_ | ϕ’) which yielded a score of improvement (Δr). Because of the nature of the phase shift fitting process described above, (r_d’_ | ϕ’) ≥ (r_d’_ | ϕ_0_) with the difference growing to a maximum when ϕ = ±180° even for data with no effect (random numbers). Therefore, we accounted for this statistical effect by running a simulation where we computed the phase shift (same process described in Figure 1e) of 1000 matrices of random numbers. For each matrix, we measured (r_d’_ | *ϕ’*) and (r_d’_ | ϕ_0_) and subtracted them as above so that we had 1000 pairs of ϕ’ and Δr. These data, along with our observed data, were fit to the function

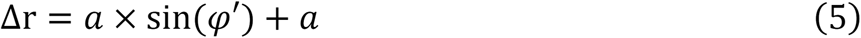

which returned *a*, the amplitude of the function. We then bootstrapped (10000 bootstrapped samples of 20 “observations” each) fits to the simulated data to obtain 95% confidence intervals for *a*. From these confidence intervals, we computed a z-score for the observed amplitude as

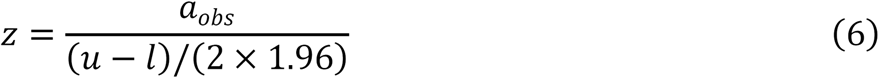

where *a_obs_* is the amplitude parameter of the fit to the observed data and *u* and *l* are the upper and lower confidence bounds from the bootstrapped fits to the simulated data, respectively.

### Diffusion Model Analysis

For binary choices, sequential-sampling models assume that upon presentation of the stimulus, the decision maker sequentially samples information from the stimulus display over time, which provides sensory evidence to a decision process. It also assumes that the decision process accumulates this evidence in a noisy manner for choosing one option over the other, here “modulation present” or “modulation absent.” Sequential-sampling models account simultaneously for choice frequency and choice response times. However, the focus here will be on choice frequencies. Let X(t) denote the random variable representing the numerical value of the accumulated evidence at time *t*. A bias, *β*, (i.e., prior beliefs about the stimulus before it is presented) can influence the initial starting position of the decision process, X(0). This initial state may either favor choice option “modulation present” (X(0) >0) or choice option “modulation absent” (X(0) <0). X(0)=0 reflects an unbiased response. (The initial states can also be given a probability distribution). The participant then samples small increments of evidence at any moment in time, which either favor response “modulation present” (dX(t) > 0) or response “modulation absent” (dX(t) < 0). The evidence is incremented according to a diffusion process. In particular, we apply a Wiener process with drift, lately called drift-diffusion model (Bogacz et al. 2006) with

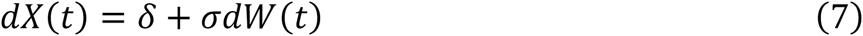

The drift rate, *δ*, describes the expected value of evidence increments per unit time. The diffusion rate, *σ,* in front of the standard Wiener process, *W(t)*, relates to the variance of the increments. Here we set *σ = 1*. The small increments of evidence sampled at any moment in time are such that they either favor response “modulation present” (*dX(t) > 0*) or response “modulation absent’’ (*dX(t) < 0*). This process continues until the magnitude of the cumulative evidence exceeds a threshold criterion, *θ*. That is, the process stops and response “modulation present” is initiated as soon as the accumulated evidence reaches a criterion value for choosing response “modulation present” (here, *X(t) = θ > 0*), or it stops and a “modulation absent” response is initiated as soon as the accumulated evidence reaches a criterion value for choosing response “modulation absent” (here, *X(t) = θ < 0*). The probability of choosing the response “modulation present” over “modulation absent” is determined by the accumulation process reaching the threshold for response “modulation present” before reaching the threshold for response “modulation absent”. The criterion is assumed to be set by the decision maker prior to the decision task. The drift rate may be related to the quality of the stimuli (i.e., the better the quality the higher the drift rate). For instance, stimuli that are easier to discriminate are reflected in a higher drift rate. In the following we consider two models. In Model 1 we assume that the physical correlation between the auditory and visual stimuli, (r_av_ | ϕ_0_), weighted by the decision maker drives the evidence accumulation process for initiating a “modulation present” or “modulation absent” response. That is, the drift rate is defined as

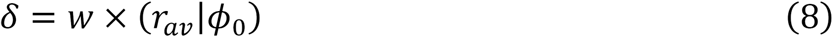

Of the 40 correlation coefficients several of them were identical (for instance, a 6Hz auditory stimulus with starting phases of +45° and −45° both resulted in a correlation of .7075) resulting in 21 unique correlation coefficients and by that in 21 different drift rates.

In Model 2 we assume that the physical correlation between the auditory and visual stimuli is distorted by a shift in phase as perceived by the decision maker. That is, the drift rate is defined by

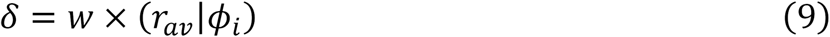

where *i* is a free parameter of the model estimated from the data and its returned value corresponds to a phase shift that is unique to each participant (*ϕ’*). The model term *ϕ_i_* relates to the initial phase term *ϕ_i_* introduced earlier and follows the same naming conventions. A phase shift unequal to 0, ±45, ±90, ±135, or ±180 results in 40 different correlation coefficients which in turn results in 40 drift rates.

### Model parameters

We assume for both models that the observed response time is the sum of the decision time, modeled by the diffusion process, and a residual time, *T_r_*, which includes the time for processes other than the decision, e.g., sensory encoding and motor components. Here, *T_r_*, is a constant for each participant. Because correlation coefficients varied between 1 and −1 but none of the participants showed perfect performances (e.g. 100% of correct responses to either a perfectly positively correlated stimulus pair or a perfectly negatively correlated stimulus pair), we allow an adjustment by including a weight for the correlations 0 ≤ *w* ≤ 1. We also allow for an a priori response bias, *β*, in favor of one response (present/absent). The decision criteria are *θ* = |−*θ*|.

In addition to these parameters, Model 2 returns a parameter *ϕ’* to account for perceived correlations based on individual phase shifts (rather than correlations based on the physical stimuli only) to be estimated from the data. To summarize: For Model 1 four parameters (*w*, *β*, *θ*, *T_r_*) are estimated from 63 data points (21 relative frequencies for correct responses, 21 mean response times for correct responses, 21 mean response times for incorrect responses. Trials with identical correlations were collapsed.) For Model 2 five parameters (*ϕ’*, *w*, *β*, *θ*, *T_r_*) are estimated from 120 data points (40 relative frequencies for correct responses, 40 mean response times for correct responses, 40 mean response times for incorrect responses).

The model was implemented in terms of the matrix approach (Adele Diederich and Busemeyer 2003) and parameters were estimated by minimizing the chi-square function (Smith and Vickers 1988),

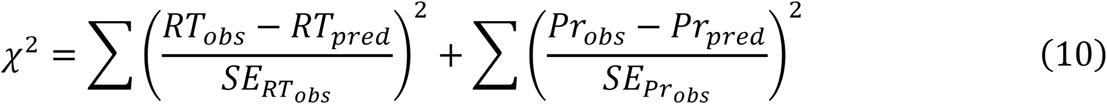

sing the optimization routine *fminsearchbnd* in MATLAB. The *fminsearchbnd* routine is similar to the standard *fminsearch* routine except that the range of the parameters of the parameters can be predetermined, for instance, positive real numbers for the residuals, or real numbers between 0 and 1 for the weights. The *fminsearch* uses the Nelder-Mead simplex search method (Lagarias et al. 1998). *SE_RT_obs__* and *SE_Pr_obs__* refer to the standard error of the observed mean response times and relative choice frequencies, respectively. Note that mean response times and relative choice frequencies are conditioned on the stimulus presented. Here we consider only the trials in which a modulation was present.

For both models, the following procedures/restrictions to parameter values were imposed in the estimation procedure: The decision criteria (absorbing boundaries) were estimated using a search grid. This was done because it quickens the estimation procedure when boundaries are integers (matrix approach). *θ* ranged from 3 to 20 in steps of 1. The residual time, *T_r_*, was restricted to 100 ms ≤ *T_r_* ≤ 800 ms and the weight to 0.0001 ≤ *w* ≤ 1. For the Model 2 parameter *ϕ_i_*, the value of *i* was restricted to integers ranging from −180 to 180 in steps of 1. For each value of *i* in Model 2, a different set of correlations were computed.

## Author Contributions

AN, RR, and MW designed the experiment. AN collected data. AN and AD analyzed the data. AN, AD, RR, and MW wrote and revised the manuscript. AN, AD, RR, and MW approved the final version of the manuscript.

## Acknowledgments

Support for this work was provided by NIH grant HD083211 to MTW and DFG DI506/15-1 to AD. We would like to thank David Simon for numerous conversations that helped guide development and analysis of the experiments and the Cognition and Cognitive Neuroscience modeling group at Vanderbilt University and especially Dr. Jeffery Annis for valuable advice during an early presentation of these data.

